# From categories to dimensions: spatio-temporal dynamics of the cerebral representations of emotion in voice

**DOI:** 10.1101/265843

**Authors:** Bruno L. Giordano, Whiting Whiting, Nikolaus Kriegeskorte, Sonja A. Kotz, Pascal Belin, Joachim Gross

## Abstract

Whether the human brain represents emotional stimuli as discrete categories or continuous dimensions is still widely debated. Here we directly contrasted the power of categorical and dimensional models at explaining behavior and cerebral activity in the context of perceived emotion in the voice. We combined functional magnetic resonance imaging (fMRI) and magneto-encephalography (MEG) to measure with high spatiotemporal precision the dynamics of cerebral activity in participants who listened to voice stimuli expressing a range of emotions. The participants also provided a detailed perceptual assessment of the stimuli. By using representational similarity analysis (RSA), we show that the participants’ perceptual representation of the stimuli was initially dominated by discrete categories and an early (<200ms) cerebral response. These responses showed significant associations between brain activity and the categorical model in the auditory cortex starting as early as 77ms. Furthermore, we observed strong associations between the arousal and valence dimensions and activity in several cortical and subcortical areas at later latencies (>500ms). Our results thus show that both categorical and dimensional models account for patterns of cerebral responses to emotions in voices but with a different timeline and detail as to how these patterns evolve from discrete categories to progressively refined continuous dimensions.

**One Sentence Summary:** Emotions expressed in the voice are instantly categorized in cortical processing and their distinct qualities are refined dimensionally only later on.

## Main text

A persistent and controversial debate in affective sciences is whether emotions are better conceptualized as discrete categories or continuous dimensions (*1, 2*). Discrete emotion theories postulate a small number of modules, each specific to a basic emotional category such as fear or anger (*3, 4*). Dimensional theories instead argue that emotions are best described along a number of continuous dimensions such as valence (reflecting the degree of pleasantness ranging from negative to positive) or arousal (reflecting the degree of intensity ranging from calm to excited) (*5, 6*).

Despite decades of continuous effort this fundamental question is still unresolved (*7*) and conflicting behavioral evidence continues to emerge both in intercultural and cross-cultural studies (*8-10*). Neuroimaging research on the cerebral bases of emotion, either felt or perceived, has not unequivocally settled this debate either (*11, 12*) and meta-analyses of large bodies of evidence can support either the notion of category-specific modules (*13*) or that of large-scale networks representing dimensional attributes (*1, 14*). Multi-voxel pattern analyses (MVPA) (*15*) have identified distributed patterns of cerebral activity allowing classification of felt or perceived emotions in others into discrete categories as well as estimation of valence and arousal dimensions (*16-24*). Yet, whether the brain represents emotional events more as discrete categories or as continuous dimensions remains unclear, in large part because the predictions of the two major theoretical positions have so far not been directly compared in a dedicated study integrating behavior with neuroimaging detailing how cerebral responses evolve in both space and time (*8, 17, 18*).

Here we address this question in humans by combining comprehensive behavioral assessments with multimodal brain-activity measurements from the same individuals at high spatial and temporal resolution. We measured cerebral activity using functional magnetic resonance imaging (fMRI) and magneto-encephalography (MEG) while participants listened to voices that densely sampled a range of perceived emotion categories and dimensional attributes (Fig. 1 and 2). This approach allowed measuring the spatiotemporal dynamics of cerebral activity during the passive perception of emotional stimuli, linking these patterns to overt behavioral responses collected after scanning and directly comparing the predictions of discrete and continuous models. We applied a multivariate analysis technique called representational similarity analysis (RSA) (*25*) to relate the perceived categorical and dimensional attributes of the stimuli (categorical and dimensional models derived from behavioral measures) to the multivariate cerebral responses (Fig. S1). With this approach, we combined multiple behavioral measures with an integrated analysis of spatial (fMRI) and spatiotemporal (MEG) cerebral activity patterns from the same participants and obtained robust converging evidence for categorical and dimensional representations of perceived vocal emotions.

**Fig. 1:**
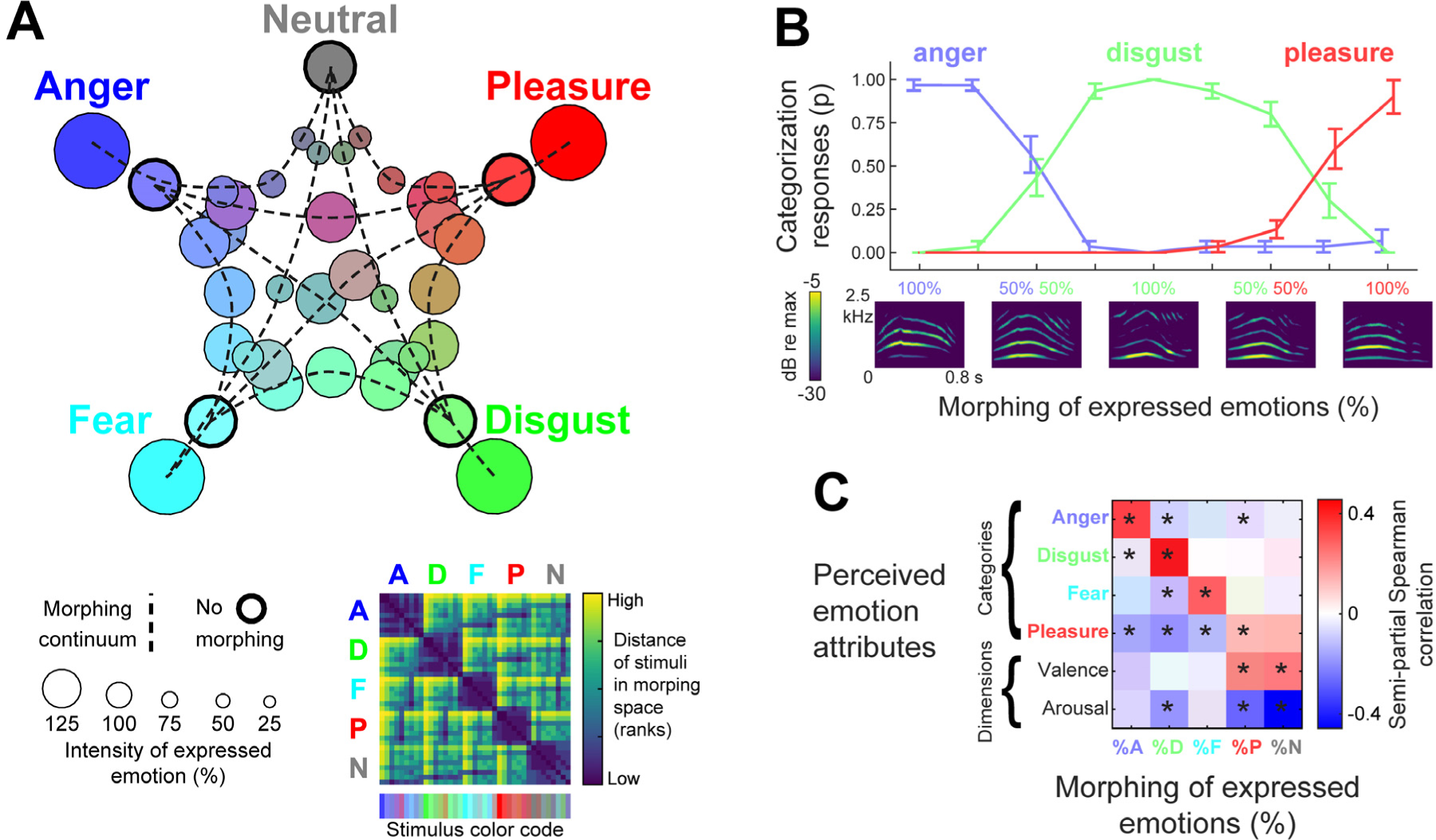
Emotions in vocal stimuli. (**A)** Auditory stimuli were generated by morphing recordings of short emotionally expressive vocalizations from the Montreal Affective Voices (*19*) portraying a neutral state and four emotions. Bottom: Euclidean distance matrix between stimuli in morphing space (each of five space dimensions measures the weight of each of the recorded vocalizations used to generate the morphed stimuli – dashed lines highlight morphing continua between pairs of recordings). Top: 2D non-metric MDS projection of the same morphing space. (**B)** Example of morphing between three vocalizations expressing different emotions (female speaker) and categorization responses (mean/error bar = bootstrap group averaged/standard error). (**C)** Effect of morphing on perceived emotion attributes: group- and speaker-averaged semi-partial correlation between morphing parameters (columns) and each of the perceived emotion attributes (rows). * p < 0.05 FWE corrected across parameters and attributes, absolute value of T(9) ≥ 4.34.

**Fig. 2:**
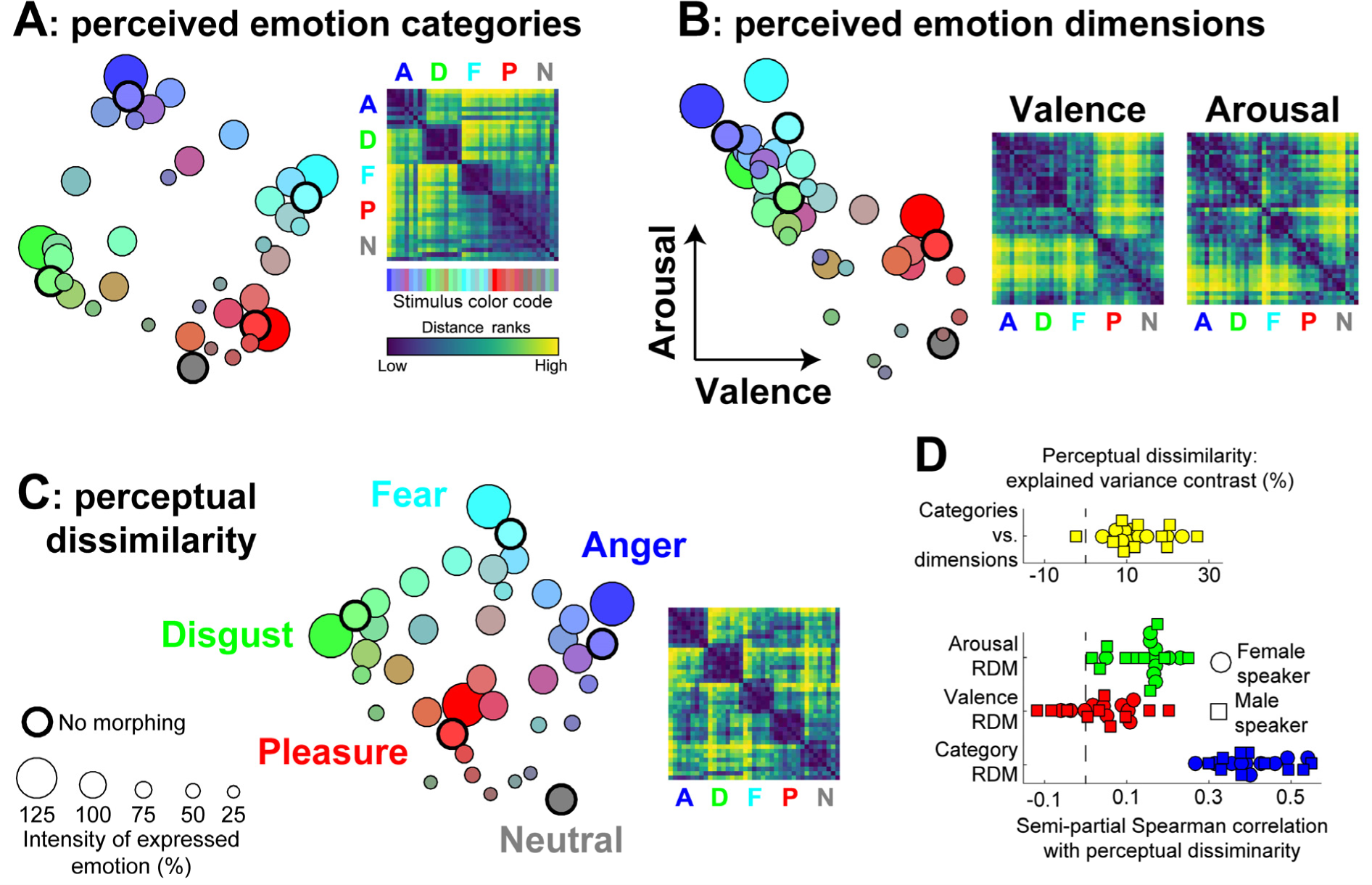
Categorical and dimensional attributes of the perceptual structure of emotions in the voice. (**A**) Right = categorical RDM averaged across participants and speakers, showing pairwise stimulus differences in emotion categorization responses for the vocal stimuli (colors measure the percent of morphing between non-morphed recordings); left = 2D MDS of categorical RDMs (non-metric INDSCAL, see Supplementary Methods). Note the strong resemblance with the perceived voice dissimilarity and MDS in C with the exception of the clustering of Pleasure and Neutral voices. (**B**) Dimensional attributes of emotions in voices: group averaged valence and arousal ratings, and corresponding RDMs and MDS representations. Note the strong valence differences between negative emotions on the one hand and Pleasure and Neutral, on the other, and the large arousal differences between Neutral and non-neutral voices. (**C**) Perceived voice dissimilarity (RDM) and corresponding 2D MDS. Note the clear clustering of each emotion and their separation from morphs with the Neutral vocalization, characterized by lower arousal (panel B). (**D**) Encoding of vocally expressed emotions in perceptual dissimilarity: bee swarm plot of semi-partial correlations of emotion attribute RDMs with the perceptual dissimilarity RDM for each experiment participant and speaker (all p < 0.05 FWE corrected across correlations and contrasts except for valence semi-partial correlation; significant T(9) ≥ 4.2), and of the percent dissimilarity variance explained by category and dimension attributes (T(9) = 10.28, p = 0.00002). See Fig. S4 for RSA analysis of the encoding of perceptual dissimilarity in fMRI and MEG response.

Auditory stimuli consisted of a homogeneous set of emotionally expressive nonverbal vocalizations obtained by morphing between the recordings of each of two actors (one female, one male) portraying four different emotions — anger, fear, disgust, and pleasure — as well as a neutral expression (*26*) while briefly uttering the vowel /a/. Morphing combined pairs of emotional vocalizations from the same actor with weights varying in 25% steps from 0 (neutral) to 125% (emotional caricature) for neutral-emotion morphs and from 0 to 100% between the four expressed emotions (Fig. 1A), resulting in 39 stimuli per actor (Audio S1-S78). Healthy participants (n=10) were each scanned in alternating sessions of fMRI and MEG (four sessions each) while performing a simple repetition detection task that ensured appropriate attention to the stimuli while avoiding an explicit focus on emotional attributes. The large amount of multimodal imaging data for each individual (8 sessions each for a total of 80 sessions) was key to adjudicating between overlapping emotion models with robust analyses. Once scanning was complete they rated the perceived dissimilarity of all (within-actor) pairs of stimuli in the absence of instructions that would bias the judgment toward a specific stimulus feature. During the last session, they evaluated the perceived emotional stimulus attributes by categorizing emotions and rating their valence and arousal (cf. Supplementary Materials and Methods).

Analysis of behavioral results confirmed that the morphing method reliably modulated perceived emotion categories and dimensions (see Fig. 1B, C for perceptual effects of morphing and Fig. 2A, B for visualization of emotion attributes in all stimuli). We quantified the relevance to perceived stimulus dissimilarity (Fig 2C) of each of three emotion-attribute distances derived from the categorization and valence and arousal ratings (emotion representational dissimilarity matrices – RDMs; Fig 2A, B; correlations between emotion RDMs ≥ .19 and ≤ 0.36, standard error of the mean–s.e.m. ≤ 0.08, T(9) ≥ 4.07, p < 0.05 family-wise error rate – FWE corrected across correlations). Although larger differences in each of the emotion attributes were associated with an increase in perceived dissimilarity (r ≥ 0.27, s.e.m. ≤ 0.03, T(9) ≥ 8.01, p < 0.05 FWE corrected across emotion RDMs), only for categories and arousal such modulation was selective, i.e. was independent of the variance shared between all emotion attributes (semi-partial correlation – s.p.r ≥ 0.15, s.e.m. ≤ 0.03, T(9) ≥ 8.98, p < 0.05 FWE corrected across emotion RDMs). Importantly, categories appeared to modulate selectively perceived dissimilarity more strongly than arousal or valence (unique explained variance contrast for categories vs. arousal or valence ≥ 29.07%, s.e.m. ≤ 2.42%, T(9) ≥ 12.52; arousal vs. valence unique explained variance contrast = 7.18%, s.e.m. = 1.58%, T(9) = 4.66; all p < 0.05 FWE corrected across contrasts) and accounted better for perceptual dissimilarity than both dimensional attributes together (percent explained variance contrast = 12.89%, s.e.m. = 1.65%, T(9) = 10.28, p = 0.00002). Thus, the behavioral data indicate that both categories and dimensions influence the perception of the emotional voice stimuli but that categories have a stronger influence.

Next, we asked where (cerebral location) and when (peri-stimulus latency) stimulus-evoked cerebral activity was significantly associated with either the categorical or the continuous emotion models. We first built fMRI RDMs reflecting at each cerebral location (voxel) the pairwise stimulus-evoked blood oxygenation level signal difference measured via fMRI, measured within a local sphere centered on that voxel (spatial fMRI searchlight = 6 mm radius). Each fMRI RDM was tested for a significant correlation with each of the three emotion-attribute RDMs (see Fig. 2A-B correlation maps and Fig. S3 and Table S3 for additional fMRI tests). We used these results to spatially constraint the subsequent MEG analysis and built MEG RDMs only at those locations that yielded significant fMRI-emotion RDM correlations (see Fig. 3A for fMRI correlation maps). The MEG RDMs were derived from pairwise stimulus-evoked magnetic signal difference at the corresponding source-space location and each peri-stimulus time point between −147 ms and 1060 ms after stimulus onset (spatiotemporal MEG searchlight = 10 mm radius, 53 ms duration and 40 ms overlap between subsequent windows; cf. Supplementary Methods and Fig. S1). The encoding of variance shared between different emotion attributes, such as the strong valence/arousal correlation apparent in Fig. 2., was teased apart from the encoding of variance unique to each of them via semi-partial correlation tests. We then contrasted the unique RDM variance explained by each of the three emotion attributes and, more importantly, by categories and both dimensional attributes together. All encoding measures generalized across the acoustical fingerprints of the male and female speakers (speaker averaged perceived emotion attributes correlated with RDMs cross-validated across speakers). Significance testing relied on a group-level permutation-based approach with cluster mass enhancement and multiple comparisons corrections across the entire analysis mask (FWE = 0.05) (cf. Supplementary Methods).

**Fig. 3:**
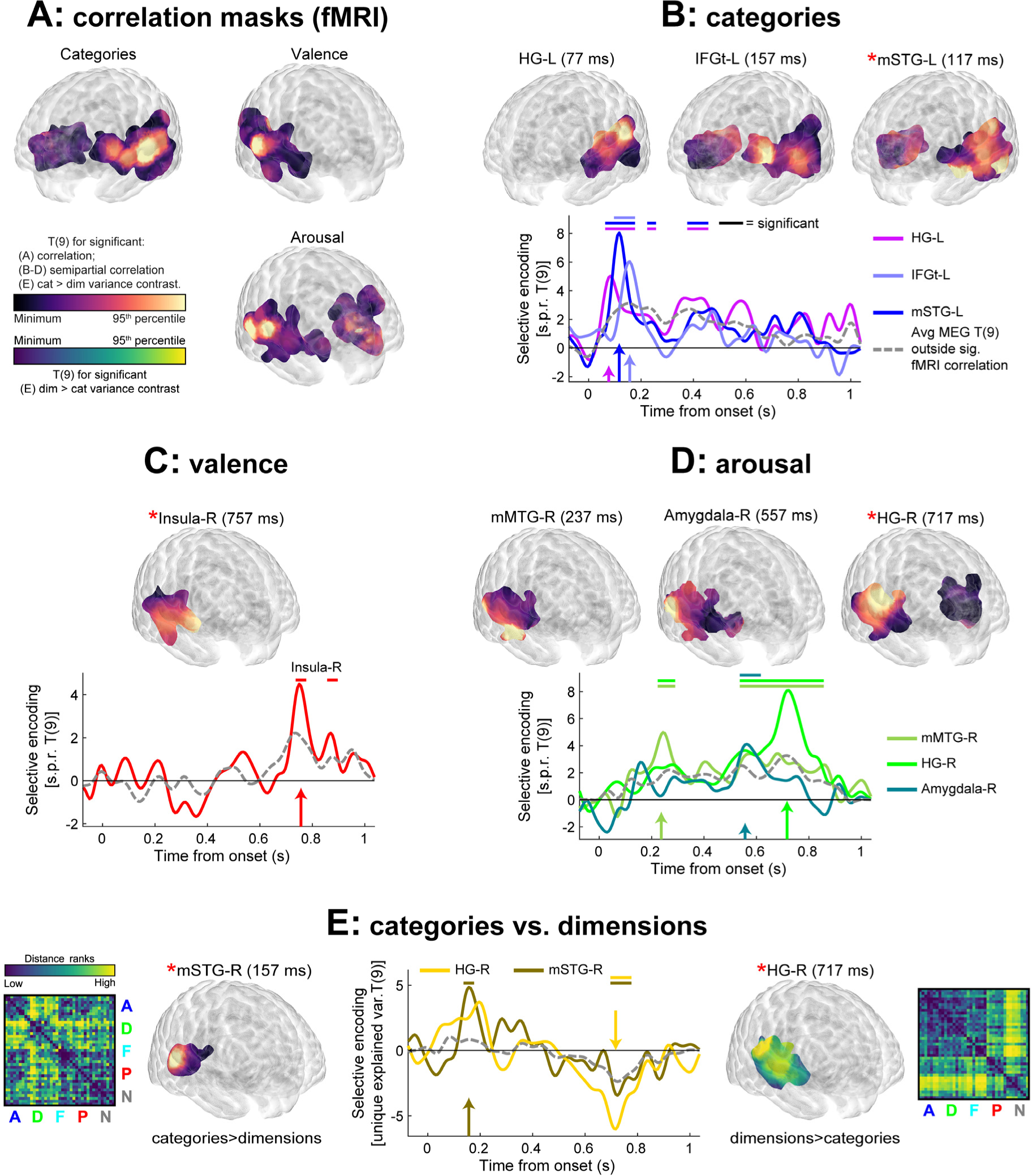
Encoding of vocally expressed emotions in the spatio-temporal cerebral network. (**A**) Non-selective encoding of perceived emotion attributes in fMRI response patterns, as measured by the correlation between emotion and fMRI response RDMs. Statistical maps thresholded for significance (p < 0.05 FWE = 0.05 corrected across voxels) are represented within a transparent MNI template (ICBM 152 2009c, see Supplementary Methods). MEG analyses were constrained spatially within the fMRI correlation masks (see Fig S3 for further fMRI results). (**B**) Selective encoding of emotion categories at selected MEG latencies (left) and in fMRI (right), as measured by the semi-partial rank correlation between category and MEG response RDMs. Statistical maps thresholded for significance (red asterisk = global spatiotemporal peak of selective-encoding statistic). The bottom panel shows the time-varying T(9) statistic used to assess the selective encoding of emotion attributes in selected ROIs. Arrows highlight the peak-effect latency for each ROI. The horizontal lines denote significant T(9) statistics – temporally overlapping significance for different ROIs emerge for shared T-statistic spatio-temporal clusters. (**C**) Selective encoding of valence and (**D**) arousal. (**E**) Contrasts of the MEG RDM variance explained uniquely by the category vs. dimension encoding model (See Fig. S3 for pairwise MEG contrasts and fMRI analyses). Also shown are the cerebral MEG RDMs at global contrast peaks. Note the similarity of the cerebral RDMs in this figure to those for the arousal and categorization behavioral judgments in Fig. 2. HG-L = left Heschl’s gyrus; m/a STG/MTG = mid/anterior superior/middle temporal gyrus; IFGt = inferior frontal gyrus, pars triangularis.

Auditory cortices bilaterally showed strong selective encoding of emotion categories from early latencies (see Fig. 3B for statistical maps and Table S1 for statistical peaks) in both primary (local MEG encoding peak at 77ms) and secondary (global MEG encoding peak at 117ms) areas of the superior temporal gyrus (STG; selective encoding extending to 517ms). At later latencies, activity patterns in these areas where characterized by selective encoding of arousal (237-837ms; global MEG encoding peak at 717ms, Fig. 3D) and, to a lesser extent, of valence in the right insula (717-877ms; global peak at 757 ms, Fig. 3C). Activity patterns selectively encoded categorical or dimensional attributes in several additional cortical and subcortical areas. The left inferior frontal gyrus (pars triangularis; IFGt) preferentially represented the set of stimuli in terms of discrete emotions from as early as 117ms after sound onset (IFGt peak at 157 ms, Fig. 3B) potentially reflecting implicit categorization processes based on feed-forward projections from the temporal cortex (*27*). Stimuli were represented in terms of their perceived arousal in the right amygdala (a subcortical structure involved in the fast detection and afferent processing of emotional signals (*28-32*)) only at relatively later latencies: starting from 237ms, (Fig. 3D), then again between 557-597ms (arousal encoding peak) and around 757ms after a brief shift of the arousal-encoding area towards orbitofrontal cortex (677ms). This temporal and differential evolution of the amygdala’s response to arousal aligns with the structure’s afferent and efferent projections to subcortical and cortical brain regions (*27, 33*).

Thus, converging evidence from three modalities—behavior, fMRI, and MEG—demonstrates that both the categorical and dimensional models explain patterns of behavioral and cerebral response to emotions in the voice—but with markedly different spatio-temporal dynamics. This may explain why previous studies have found evidence in support of either one or the other model (*13, 14, 16-24*). Our results shed significant light onto the debate by showing that categorical and dimensional representations unfold along different timelines in different cerebral regions, adding a much-needed temporal dimension to the picture of cerebral processing of perceived emotion that so far has remained rather static. We find that the amygdala showed strong associations with the arousal dimension at latencies within 237-757ms. This is consistent with previous findings of selective impairments of arousal, but not valence recognition in amygdala lesions (*34, 35*) and with neuroimaging of healthy individuals showing representation of arousal but not valence in the amygdala (*36*). In contrast, the valence dimension was weakly associated with perceptual representations and was represented in the brain only at later latencies (>700ms in the insula). Overall, the selective encoding of dimensional attributes in the amygdala and insula is in agreement with the involvement of a “salience” network (*11*) linking the processing of emotional states and events across species (*37, 38*) and thought to represent a phylogenetic precursor for communicative behavior in primates and humans (*20, 39, 40*). In other cerebral areas, however, stimulus representations appeared to evolve in time from one model to the other, such as right auditory cortex that represents stimuli first in terms of their categorical structure at early latencies and then in terms of their perceived arousal at later latencies, subsequent to their initial encoding in subcortical structures (Fig. 3E). The representational dynamics observed for the right auditory cortex thus suggests a transition from an early dominance of feed-forward sensory processing to late attentional modulations resulting from feedback signals transmitted through lateral and medial cortical connections from the amygdala (*39, 41*).

Finally, we perfomed a direct comparison of the categorical and continuous models by asking when and where patterns of neural activity reflected one theoretical account more than the other. For this, we initially calculated the contrast of RDM variance explained uniquely by each of the emotion attributes and then contrasted the explanatory power of the categorical and dimensional models (see Fig. 3E and S3 for contrast maps and Tables S1 and S2 for statistical peaks). The categorical model uniquely explained significantly more MEG RDM variance than either valence or arousal or both combined at early latencies (157ms) in the right auditory cortex centered on mid-STG (mSTG; categories vs. valence contrast significant also at 197 and 357ms). Conversely, the dimensional model uniquely explained significantly more MEG RDM variance at later latencies (717-757 ms) in a similar area of the right auditory cortex (arousal vs. categories contrast significant at 717ms; arousal vs. valence contrast significant at 637-677 ms).

In summary, by enabling a direct contrast of the predictions of the two models, our results provide crucial insight into the category vs. dimension debate. Statistical comparison of the predictions of the two models yielded unequivocal evidence for a clear prevalence of the categorical model: the perceptual structure of the stimuli was more related to categories than dimensions and spatiotemporal activity patterns in widespread areas of the auditory cortices were associated from early latencies on (as early as 77ms post-onset) with the categorical stimulus structure. The contrast of variance uniquely explained by categories and dimensions was significant in the right auditory cortex around 157ms after stimulus onset. However, dimensional representations become more prevalent at later latencies in the auditory cortex, subcortical areas, and orbitofrontal cortex, suggesting progressive refinement of emotional stimulus representations from formation of main emotional categories well suited to trigger fast adaptive reactions to increasingly fined-grained representations modulated by valence and arousal. Overall, our results provide a comprehensive characterization of the spatiotemporal dynamics of perceived emotion processing by the brain and demonstrate how both categories and dimensions are interwoven into rich and complex representations initially dominated by categories then progressively refined into dimensions.

## Acknowledgments

Supported by UK’s Biotechnology and Biological Sciences Research Council (grants BB/M009742/1 to JG, BLG, SAK, and PB, and BB/L023288/1 to PB and JG), by the French Fondation pour la Recherche Médicale (grant AJE201214 to PB), and by Research supported by grants ANR-16-CONV-0002 (ILCB), ANR-11-LABX-0036 (BLRI), and the Excellence Initiative of Aix-Marseille University (A*MIDEX). Conceptualization: BLG, PB; Methodology: BLG, CW, NK, SAK, PB, JG; Software: BLG; Validation: BLG; Formal Analysis: BLG, CW, JG; Investigation: BLG, CW; Resources: BLG, PB; Data Curation: BLG, CW; Writing - Original Draft: BLG, CW, SAK, PB, JG; Writing – Review & Editing: BLG, CW, NK, SAK, PB, JG; Visualization: BLG; Supervision: BLG, PB, JG; Project Administration: JG; Funding Acquisition: BLG, SAK, PB, JG. We thank Dr. Olivier Coulon and Dr. Oliver Garrod for help with the development of the 3D glass brain.

## Supplementary Materials and Methods

### Participants

Ten right-handed healthy adults (5 female; age from 19 to 38, mean = 25.1) participated in this study. All participants had normal hearing as assessed by an audiogram, provided written informed consent, and received financial compensation of £6/hour for their participation. The study was conducted in accordance with the Declaration of Helsinki and was approved by the local ethics committee (College of Science and Engineering, University of Glasgow).

### Stimulus material

Stimuli consisted of nonverbal emotionally expressive vocalizations from the Montreal Affective Voices database (*26*) and were produced by two actors (one male, one female). Each actor produced five vocalizations (vowel /a/) expressing: anger, disgust, fear, pleasure, and neutral. Vocalizations normalized in root mean square (r.m.s.) amplitude were then used to generate the stimulus set by morphing between each pair of vocalizations from the same speaker.

Voice morphing was performed using STRAIGHT (*42*) in Matlab (Mathworks, Inc, Natick, USA). STRAIGHT performs an instantaneous pitch-adaptive spectral smoothing in each stimulus for separation of contributions to the voice signal arising from the glottal source versus supralaryngeal filtering. A voice stimulus is decomposed by STRAIGHT into five parameters (f0, frequency, duration, spectrotemporal density, and aperiodicity) that can be manipulated and combined across stimuli independently of one another. Time-frequency landmarks to be put in correspondence across voices during morphing were manually identified in each stimulus, and corresponded to the frequencies of the first three formants at onset and offset of phonation. Morphed stimuli were then generated by resynthesis based on the linear (time and aperiodicity) and logarithmic (f0, the frequency structure and spectrotemporal density) interpolation of these time-frequency landmarks.

Two types of morphing continua were produced: 1) between neutral and each of the four emotions (neutral-anger, neutral-disgust, neutral-fear, and neutral-pleasure), and 2) between pairs of emotions (anger-disgust, anger-fear, anger-pleasure, disgust-fear, disgust-pleasure, and fear-pleasure). The morphing continuum between neutral and each emotion consisted of 6 stimuli, progressing in acoustically equal steps of 25% (e.g., neutral 100% → neutral 75%/anger 25% → neutral 50%/anger 50% → neutral 25%/anger 75% → anger 100% → anger 125%). The 125% emotion was generated by extrapolating along the neutral-emotion dimension to create a caricatured emotion. The morphing continuum between pairs of emotions consisted of 5 stimuli, again progressing in acoustically equal steps of 25%. In total, 78 stimuli were used in the experiment, consisting of 39 stimuli for each speaker (cf. Audio Files S1-S78). They were normalized to the average duration of 796 ms using pitch-preserving time-stretching algorithms, and then in root mean square amplitude.

### Experimental design

Each individual took part in 11 experimental sessions. Neuroimaging data were collected during the first 8 sessions (4 fMRI and 4 MEG; imaging modalities alternated with fMRI first for half of the participants; MEG at least 3 days after prior fMRI session to avoid magnetization artefacts). Behavioral data were collected during the last three sessions, perceived categorical and dimensional emotion attributes being estimated only during the last session to avoid biases towards either during the rest of the experiment.

On each run of the fMRI and MEG acquisition (20 runs per fMRI session and at least 78 runs per participant across all of the MEG sessions), participants were presented with all of the stimuli from one speaker (random speakers order on each pair of subsequent blocks; inter-stimulus interval – ISI – jittered between 3 and 5 s) while carrying out a one-back repetition detection task (1 repetition per run; random selection of repeated stimulus; group averaged p correct = 98%; s.e.m. = 0.2%).

Throughout the session, participants were instructed to fixate a cross, presented in greyscale (screen field of view = 19×80 and 26×19 degrees for fMRI and MEG, respectively).

At each of the first two behavioral sessions, participants rated the dissimilarity between all of the stimuli from the same speaker (speaker order counterbalanced across participants). On each trial, they were presented with one of the possible 741 pairs of sounds (within-pair ISI = 250ms; random within-pair order) and were asked to rate how dissimilar they were by placing a slider along a visual analogue scale marked “very similar” and “very dissimilar” at the two extremes. They could listen to the pair of stimuli as many times as necessary before giving a response. This experimental phase was preceded by an initial familiarization phase during which participants were presented with all of the sound stimuli two times (ISI = 250 ms; random order). In this phase, they were instructed to estimate the maximum and minimum between-sound dissimilarity, so as to optimize the usage of the rating scale in the subsequent experimental phase. The procedure was initially practiced with a set of 10 vocalizations not included in the main experiment.

During the last behavioral session, participants performed two tasks – emotion categorization and ratings of dimension attributes. In the categorization task, they identified the emotion as being anger, disgust, fear, or pleasure. In the rating tasks, participants rated each stimulus on arousal (low to high) and valence (negative to positive) using an on-screen slider. Before the experiment began, participants were given 10 practice trials for both the categorization and rating tasks on an independent set of vocal stimuli. Participants were then familiarized to the entire stimulus set before the first block. On each block of trials, participants carried out either the category or rating tasks (alternated across blocks) for all of the stimuli from the same speaker (pseudo-random order of speaker gender with not more than two subsequent same-gender blocks). Throughout the session, each of the two tasks was repeated three times for each of the speakers, for a total of 12 blocks of trials.

Sound stimuli (sampling rate = 48 kHz; bit depth = 16 bit) were presented through electrostatic headphones (NordicNeuroLab, Bergen, Norway) for fMRI, Etymotic ER-30 tubephone for MEG, and during the behavioral sessions through BeyerDynamic DT 770 Pro headphones receiving the audio signal from the Audiophile 2496 sound card amplified with a Mackie 1604-VLZ PRO monitor system. The MEG tubephone system introduced strong spectral coloring of the sound stimuli and suppressed heavily frequencies > 6 kHz. Stimuli for all sessions were consequently low-pass filtered at 5 kHz. Flat-frequency response for the MEG audio stimulation chain was achieved through inverse filtering methods.

### Neuroimaging data acquisition

fMRI scans were acquired with a Siemens 3T Trio scanner, using a 32-channel head coil. Functional multiband echo planar imaging (EPI) volumes were collected with a repetition time (TR) of 1s (echo time TE = 26 ms; flip angle = 60; multiband factor = 4; GRAPPA = 2). Each functional volume included 56 slices of 2.5 mm thickness (inter-slice gap = 2.5 mm; interleaved even acquisition order (interleaved even) in an axial orientation along the direction of the temporal lobe, providing nearly whole-brain coverage. The in-plane voxel size was 2.5 mm^2^ (78 × 78 matrix). A whole-brain, high-resolution, structural T1-weighted MP-RAGE image (192 sagittal slices, 256 × 256 matrix size, 1 mm^3^ voxel size) was also acquired to characterize the subjects’ anatomy. In each of the fMRI sessions, we also collected a field map to correct for geometric distortions in the EPI volumes caused by magnetic field inhomogeneities (*43*).

MEG recordings were acquired with a 248-magnetometers whole-head MEG system (MAGNES 3600 WH, 4-D Neuroimaging) at a sampling rate of 1017.25 Hz. Participants were seated upright. The position of five coils, marking fiducial landmarks on the head of the participants, was acquired at the beginning and at the end of each block.

### Analysis of behavioral data

We initially assessed the effect of voice morphing on perceived emotion attributes (five morphing % parameters describing each experimental stimulus – one for each expressed emotion and one for the neutral vocalization; six measures of perceived emotion – four emotion categorization probabilities plus valence and arousal ratings). The five morphing parameters were not orthogonal because for each stimulus only two at best had a non-zero value (average Spearman correlation between morph parameters = −0.24; STD = 0.02). For this reason, the perceptual effect of morph parameters was assessed independently of their shared variance by measuring their Spearman semi-partial correlation (s.p.r) with the measures of perceived emotion. Significance testing for all of the analyses in this study relied on a permutation-based group-level random effects (RFX) approach. Here, we: [1] estimated independently for each participant and speaker the null s.p.r distribution for each of the 30 morph/emotion pairs by permuting randomly the stimulus labels (N permutations = 100,000; same permutations across speakers and morph/emotion pairs, but not across participants); [2] averaged across speaker genders permuted and unpermuted s.p.r. converted to the Fisher Z scale; [3] subtracted the median bias of the null s.p.r distributions from both the permuted and unpermuted s.p.r.; [4] computed the T(9) test for the group-average permuted and unpermuted s.p.r.; [5] finally established significance thresholds for the unpermuted T(9) tests as the 95^th^ percentile of the distribution of the maximum of the absolute value of the permuted T(9) statistics (two-sided inference) across pairs of morph parameters with perceived emotion measures, thus controlling for family-wise error (FWE) at a 0.05 level.

Subsequent modeling of behavioral data assessed the association of perceived stimulus dissimilarity (pairwise dissimilarity ratings) with perceived emotion categories, valence and arousal (each transformed to a pairwise stimulus distance – emotion representational dissimilarity matrix RDM). The valence and arousal RDMs measured the absolute pairwise difference in valence and arousal ratings, respectively. The category RDM was defined as the Euclidean distance between stimulus-specific categorization response profiles (e.g., categorization profile consisting of 10, 2, 1 and 0 anger, disgust, fear and pleasure responses). All emotion RDMs were computed independently for each participant and speaker. Significance testing relied on a similar approach as for the analysis of the effect of morph parameters on perceived emotion attributes (N permutations = 100,000; reshuffling of rows and columns of distance matrices). Importantly, however, we opted for one-tailed inference to assess the significance of the correlation between emotion RDMs themselves (FWE = 0.05 across the three pairwise correlations) and the correlation and semi-partial correlations between emotion RDMs, on the one hand, and the dissimilarity ratings, on the other. Additional contrasts compared the proportion of dissimilarity rating variance uniquely explained by each of the three emotion RDMs, and by the category RDM vs. the dimensional attribute RDMs together (square root of unique explained variances Fisher Z transformed prior to contrast; two-tailed inference for all contrasts; FWE = 0.05 adjusted across the three pairwise contrasts; two-tailed percentile p-values for categories vs. dimensions contrast).

### Preprocessing of neuroimaging data

Analyses were carried out in Matlab using SPM12, Fieldtrip (*44*), GLMdenoise (*45*) and custom code. The initial preprocessing of fMRI and MEG data produced for each participant the stimulus-specific responses further analyzed to assess the encoding of emotion attributes (see below). Functional MRI images from all runs were realigned to the first image in the first run and unwarped to correct for movement-by-distortion interactions (full width at half maximum – FWHM = 5 and 4 mm for realignment and unwarp, respectively; for both 7th degree B-spline for interpolation), and slice time corrected to the onset of the temporally central slice. Anatomical volumes were co-registered to the grand-average of the preprocessed functional volumes and segmented into grey matter, white matter, and cerebro-spinal fluid. Diffeomorphic Anatomical Registration using Exponentiated Lie algebra (*46*) (DARTEL) was used to create a common brain template for all of the participants. An initial group DARTEL grey-matter mask was created by considering all non-cerebellum voxels with a grey-matter probability > 0.1. The final analysis mask for each individual was given by the 6-connected voxels within the conjunction of the group mask deformed to native space with the voxels associated with a participant-specific grey-matter probability > 0.25.

For each participant, the 80 fMRI runs (40 for each of the two speaker genders) were divided into 5 mixed-gender groups of 16 runs each (interleaved assignment of runs to groups). Unsmoothed native-space data within the analysis mask for each group of runs were analyzed within a massively univariate general linear model (GLM) that estimated the fMRI response specific to each stimulus. Stimulus-specific regressors were created by convolving a sound on-off binary time-series with the canonical hemodynamic response function (HRF). The GLM included a high-pass discrete cosine transform (DCT) filter (cut off = 128 s), the head motion regressors estimated during the realignment step and a run-specific intercept. The GLM also included additional noise regressors that modeled temporal effects unrelated to the stimulus condition (e.g., blood pulse). They were estimated independently for each of the groups of runs and participants using GLMdenoise ((*45*); default polynomial detrending replaced with DCT filter), resulting in N noise regressors = 6 on average (across-participant STD = 2).

Several initial steps of the preprocessing of MEG data were carried out on the unsegmented data from each run. Infrequent SQUID jumps (observed in 2.3% of the channels, on average) were repaired using piecewise cubic polynomial interpolation. For each participant independently, we then removed channels that consistently deviated from the median spectrum (shared variance < 25%) on at least 25% of the runs (N removed channels = 8.4 on average; STD = 2.2). Runs associated with excessive head movements or MEG channels noise or containing reference channel jumps were finally discarded, leaving on average 75.9 runs per participant (range = 65– 84; average maximum coil movement across blocks and participants = 5 mm; STD = 1 mm). Environmental magnetic noise was removed an initial time using regression based on principal components of reference channels. Both the MEG and reference data were then filtered using a forward-reverse 70 Hz FIR low-pass (−40 dB at 72.5 Hz), a 0.2 Hz elliptic high-pass (−40 dB at 0.1 Hz) and a 50 Hz FIR notch filter (−40 dB at 50 ± 1Hz), and were subsequently resampled to 150 Hz. Residual magnetic noise was then removed applying once more the same method as for the full-resolution signal. ECG and EOG artifacts were removed using ICA (runica on 30 components) and were identified based on the time course and topography of IC components (*47*). MEG data from each run was finally segmented into trials (−0.2 to 1.3 s after sound onset).

A native-space source-projection grid with a resolution of 3.5 mm was prepared for each participant by resampling the native-space analysis mask for the fMRI data. Depth-normalized lead fields were computed based on a single shell conductor model. Source-projection filters were then computed for each run using LCMV beamformers (regularization = 5%; sensor covariance across all trials excluding repetitions) and reduced to the maximum-variance orientation across all runs. Source-projected stimulus-specific time courses were finally averaged within 5 independent mixed-gender groups of runs (interleaved assignment of runs to groups), leading to a reduction of the computational burden for subsequent data-analysis steps.

### Cerebral encoding analysis

We implemented a whole-brain searchlight representational similarity analysis (RSA; (*25*); Fig. S1) to assess the encoding of perceived emotion attributes in multivariate spatial (fMRI) and spatiotemporal (MEG) cerebral response patterns. We followed the same approach adopted for the analysis dissimilarity rating data, and measured here the association between emotion and cerebral response RDMs.

For fMRI, cerebral RDMs were computed in native space within a spherical region (6 mm diameter) centered at each grey-matter mask location (at least 50% in-mask voxels). In particular, we computed the cross-validated Mahalanobis distance between stimulus-specific response patterns (Mahalanobis whitening of stimulus-specific GLM estimates using the GLM residuals within the searchlight) by cross-validating the response pattern covariance across the 5 groups of mixed-gender runs, and finally converting it to a (whitened) Euclidean distance (*48, 49*). For MEG, cerebral RDMs were computed within a spatiotemporal searchlight of 10 mm diameter and 50 ms temporal window from −0.15 to 1.1 seconds from onset with 15 ms of overlap between subsequent temporal windows. For each searchlight, we derived the cross-validated Euclidean distance between stimulus-specific beamformed time-courses from the covariance between stimulus-specific response patterns cross-validated between the 5 groups of mixed-gender runs.

RSA analyses assessed, in order: [1] the Spearman correlation between cerebral and emotion RDMs (non-selective encoding; one sided inference); [2] the Spearman s.p.r between cerebral and emotion RDMs (selective encoding; one sided inference); [3] the pairwise contrasts of the unique cerebral RDM variance explained by each of three pairs of emotion RDMs, and [4] the explained cerebral RDM variance contrasts between the categorical and dimensional models (valence and arousal together; two-sided inference for all contrasts). Importantly, within each imaging modality, we: [1] computed all encoding measures in native space and carried out group-level RFX inference (T tests) on the encoding maps transformed to the group DARTEL space (FWHM of Gaussian smoothing of native-space encoding maps = 8 for both fMRI and MEG); [2] used cluster mass enhancement of the group-level statistics, permutations included (permutation of rows and columns of RDMs, as for analysis of perceptual dissimilarity; 3D and 4D spatiotemporal cluster mass enhancement for fMRI and MEG, respectively; cluster-forming threshold of T(9) = 1.83 and 2.26 for one- and two-sided inference, respectively; (*50*)); [3] mitigated the multiple comparison problem by constraining analysis masks at each testing step within the significance mask from the previous step (correlations tested at whole brain and latency-range levels; s.p.r within significant correlation masks; variance contrasts within significant s.p.r masks). For MEG we assessed the initial cerebral response/emotion attribute RDM correlations within the significant correlation masks from the fMRI analysis, providing a further mitigation of the multiple comparison problem and capitalizing on the superior spatial and temporal specificity of fMRI and MEG, respectively. At all steps, we corrected for multiple comparisons within the entire analysis mask by establishing significance thresholds for the non-permuted cluster-mass enhanced T statistics as the 95^th^ percentile of the permutation distribution for within-mask CM enhanced maxima for one-sided tests and as the 2.5^th^ and 97.5^th^ percentiles of within-mask minima and maxima, respectively for two-sided inference (maximum-statistic approach; FWE = 0.05).

### Visualization

The non-metric MDS models in Fig. 1 and 2 were computed using the R-package SMACOF (*51*). We modeled the dissimilarity ratings and the emotion RDMs using an inter-individual difference scaling model (INDSCAL), avoiding well known distortions of the representational geometry associated with group averaging of distance data (*52*). The 3D glass brains in Fig. 3 and S3 comprised two components: [1] a mesh of the ICBM 152 2009c Nonlinear Asymmetric template (*53*); [2] the functional blobs, rendered by first modeling the surface of each blob with a 3D mesh, and then projecting onto it the volumetric statistical map it circumscribed (maximum projection within 7 mm radius sphere centered at mesh vertex). All meshes and projections were computed within BrainVISA (http://brainvisa.info/), and were rendered using a custom OpenGL shader for the transparency effect.

---

**Fig. S1:**
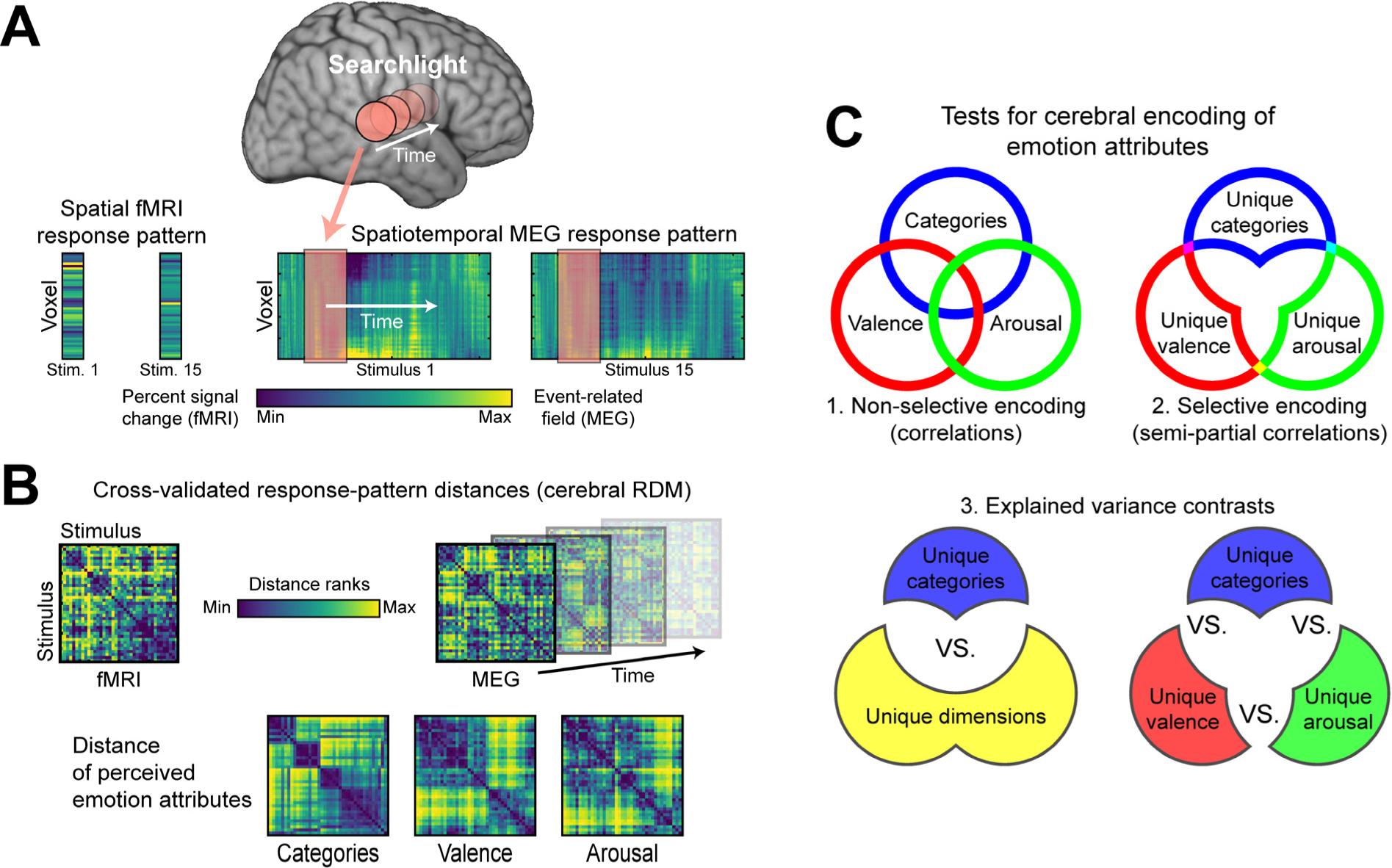
Representational similarity analysis of the encoding of perceived emotion attributes in spatio-temporal cerebral response patterns. (**A**) Spatial (fMRI) and spatio-temporal (MEG) stimulus-specific response patterns (percent signal change and source-space event-related fields for fMRI and MEG, respectively) were extracted within a grey-matter spherical searchlight (radius: 6 and 10 mm for fMRI and MEG, respectively). The MEG spatio-temporal searchlight had a duration of 50ms and an overlap with subsequent windows of 15ms. (**B**) Cerebral encoding analyses involved the measurement of the association between the pairwise distance of stimulus-specific cerebral responses (cerebral representational dissimilarity matrices – RDMs) and the pairwise distance of stimulus-specific perceived emotion attributes (emotion RDMs). (**C**) For each modality, encoding tests were carried out in subsequent steps within nested significance masks. First, we assessed for the non-selective encoding of each emotion attribute by measuring the correlation of each with the cerebral RDMs. Note that emotion attribute RDMs are correlated (see main text) and explain overlapping portions of the cerebral RDM variance. The second step focused on portions of the cerebral RDM variance explained uniquely by each emotion attribute, and assessed their selective encoding by means of their semi-partial correlations with the cerebral RDMs. Finally, we directly contrasted the explained cerebral RDM variance specific to the category- or dimension-encoding model, and the unique explained cerebral RDM variance specific to each of the three perceived emotion attributes (three pairwise contrasts). Subsequent tests were carried out independently with the two modalities, with the important exception of constraining the non-selective MEG encoding tests within those brain areas characterized by non-selective emotion attribute encoding in fMRI data.

**Fig. S2:**
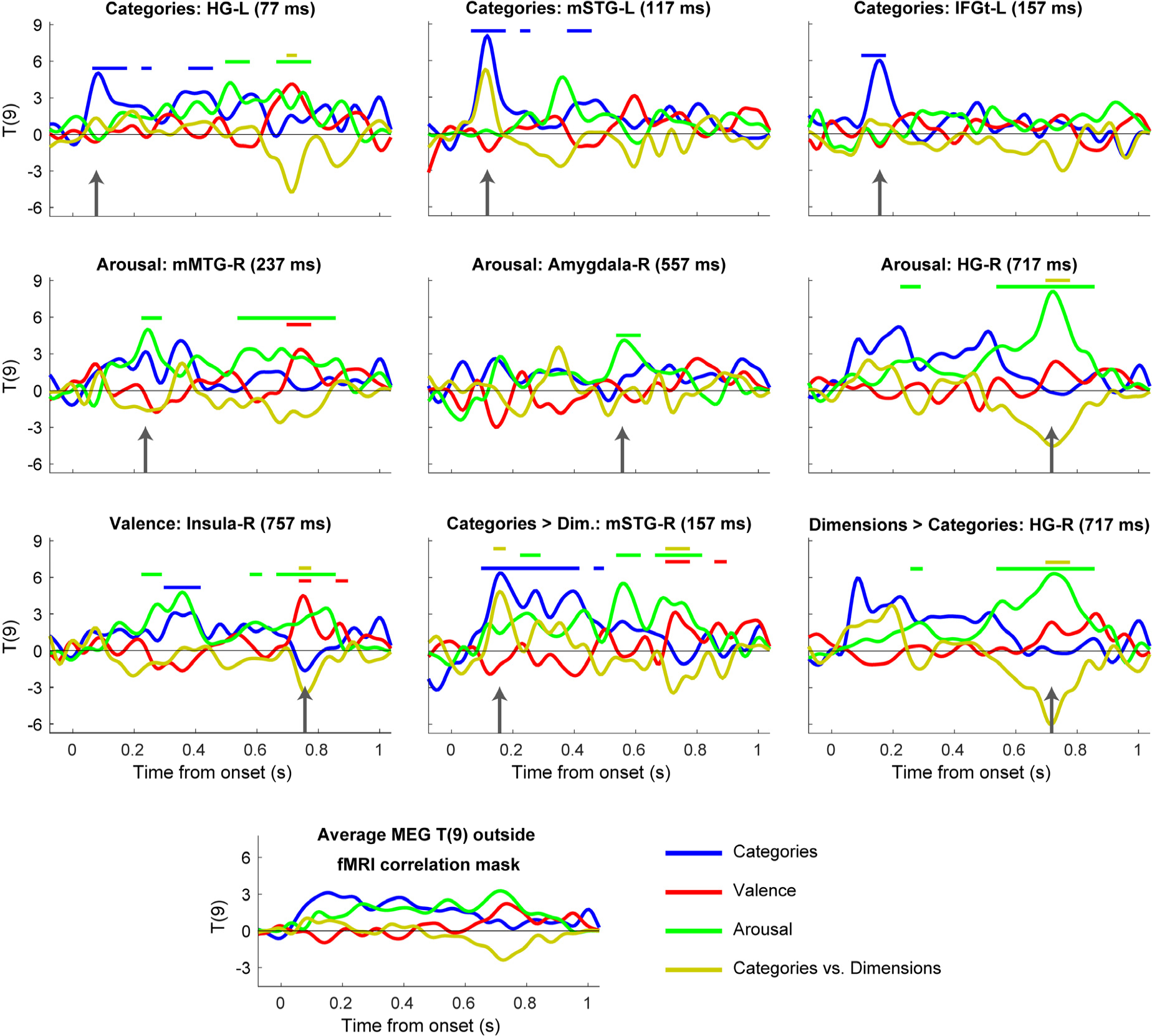
Time-varying emotion encoding in MEG data. Time courses for the MEG T(9) statistics used to assess the selective encoding of emotion attributes and to contrast the MEG RDM variance explained by the category vs. dimension models. The time courses are extracted from the global and local peaks of the MEG statistical maps (see Table S1). HG-L = left Heschl’s gyrus; m STG/MTG = mid superior/middle temporal gyrus; IFGt = inferior frontal gyrus, pars triangularis. See Fig. 3 for more details.

**Fig. S3:**
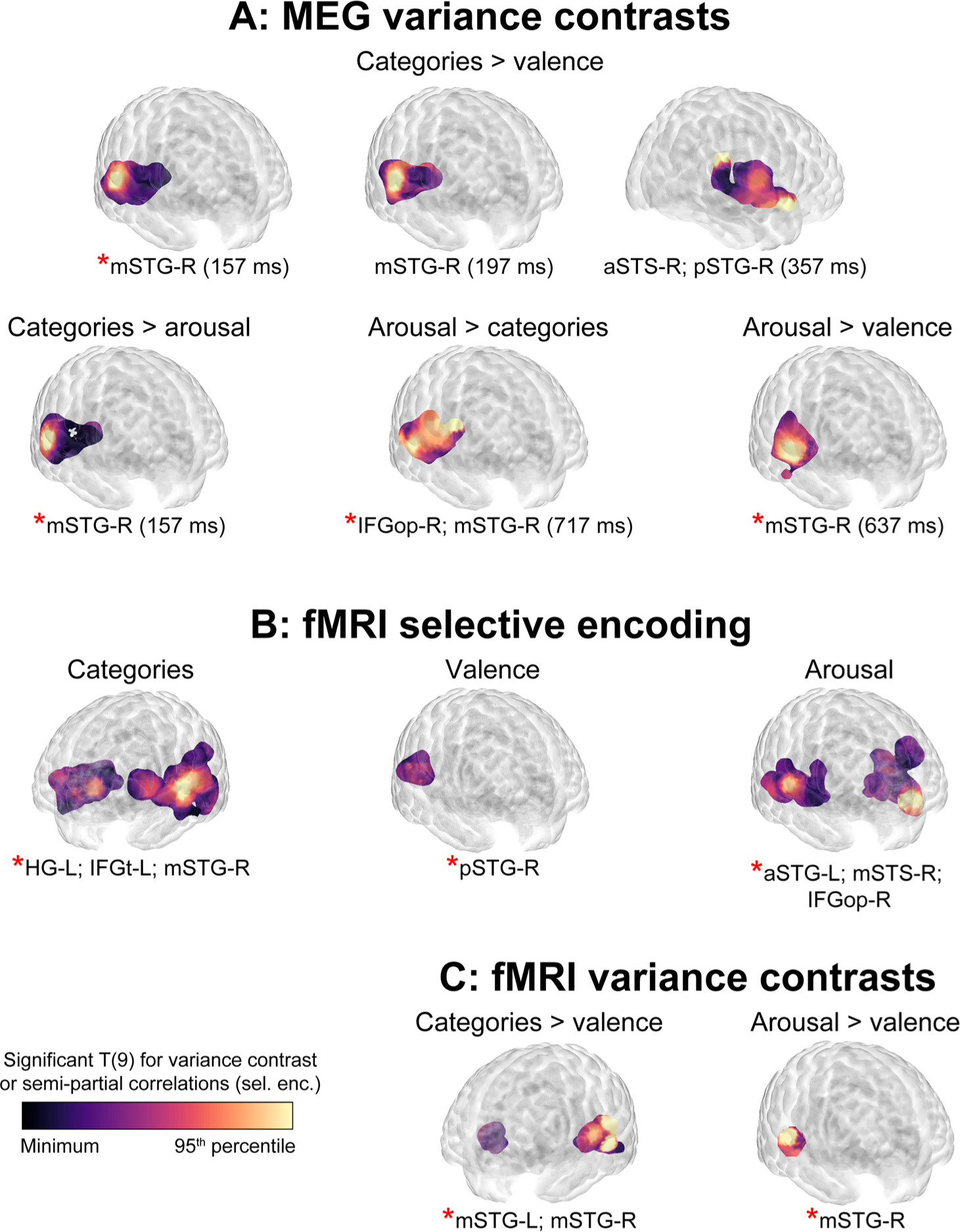
Encoding of the perceived emotional attributes of voices in spatiotemporal (MEG) and spatial (fMRI) cerebral response patterns. All effects significant at p < 0.05 FWE corrected across voxels (fMRI and MEG) and time points (MEG) within the analysis masks. p/m/a STG-R = right posterior/mid/anterior superior temporal gyrus; STS = superior temporal sulcus; IFGop/t = inferior frontal gyrus, pars opercularis/triangularis; HG = Heschl’s gyrus. See Fig. 3 and Tables S2 and S3 for more details.

**Fig S4:**
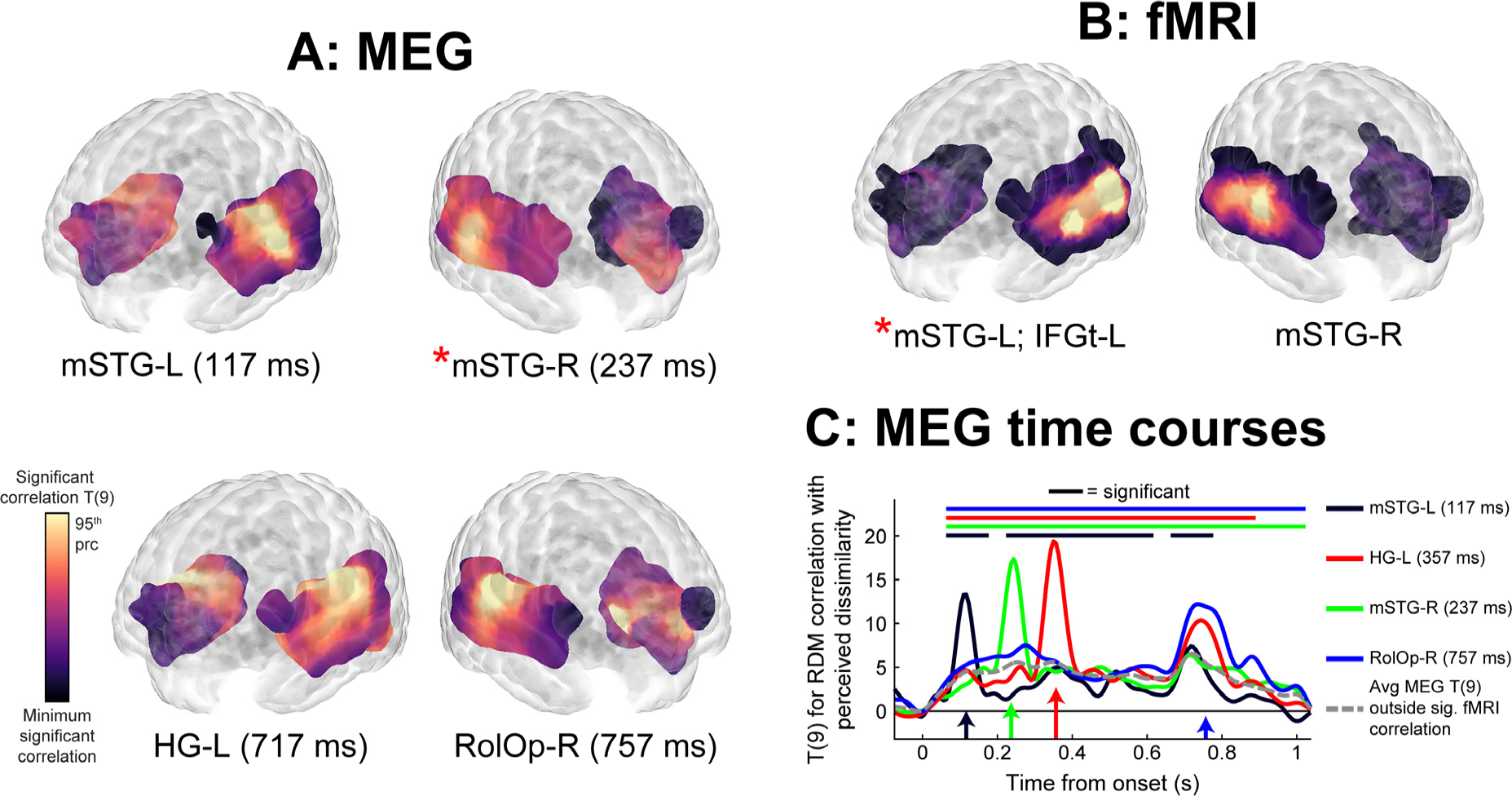
Encoding of perceived dissimilarity in cerebral response patterns. We tested for the cerebral encoding of the perceived dissimilarity of the emotionally expressive stimuli using the same correlation approach as for the emotion attribute RDMs. The MEG analysis was constrained within the mask for a significant correlation between fMRI RDMs and dissimilarity ratings data. All effects shown significant at FWE < 0.05. mSTG-L = left mid superior temporal gyrus; IFGt = inferior frontal gyrus, pars triangularis; HG = Heschl’s gyrus; RolOp = Rolandic operculum. See Figure 3 for more details.

**Table S1:** **Global and local peaks of main emotion encoding analyses in MEG data.** The table lists global and local peaks in the GLM T-maps. Anatomical labels are based on the Automated Anatomical Labeling (AAL) atlas. HG-L = left Heschl’s gyrus; mSTG/MTG = mid superior/middle temporal gyrus; IFGt = inferior frontal gyrus, pars triangularis; Ins = insula; Amy = amygdala; * = global peak; effect size = group averaged of native-space semi-partial correlations at corresponding voxels and of explained variance contrasts; SEM = standard error of the participant average. All effects significant at p < 0.05 FWE corrected across voxels and time points.

| **Effect** | **Anatomical label** | **Latency (ms)** | **MNI coordinates** |  |  | **T (9)** | **Effect size (s.e.m.)** |
| --- | --- | --- | --- | --- | --- | --- | --- |
| Categories | HG-L | 77 | -62 | -30 | 20 | 4.93 | 0.11(0.02) |
| Categories | mSTG-L* | 117 | -42 | -2 | -18 | 8.19 | 0.09(0.01) |
| Categories | IFGt-L | 157 | -49 | 23 | -3 | 6.55 | 0.09(0.01) |
| Valence | Ins-R* | 757 | 41 | -2 | -8 | 5.1 | 0.15(0.02) |
| Arousal | mMTG-R* | 237 | 56 | -17 | -18 | 5.09 | 0.14(0.02) |
| Arousal | HG-R | 717 | 58 | -10 | 17 | 8.04 | 0.26(0.03) |
| Arousal | Amygdala-R | 557 | 14 | -2 | -8 | 4.2 | 0.11(0.02) |
| Cat>Dim | mSTG-R* | 157 | 66 | -37 | 4 | 5.31 | 1.47(0.04) |
| Dim>Cat | HG-R* | 717 | 64 | -17 | 27 | 6.12 | 7.10(0.10) |

**Table S2:** **Global and local peaks of pairwise contrasts between the MEG variance explained by each emotion attribute.** The table lists global and local peaks in the GLM T-maps. Cat = categories; Val = valence; Aro = Arousal; m/pSTG-R = right mid/posterior superior temporal gyrus; aSTS = anterior superior temporal sulcus; IFGop = left inferior frontal gyrus, pars opercularis. All effects significant at p < 0.05 FWE corrected within the analysis mask. See Table S1 for further details.

| **Effect** | **Anatomical label** | **Latency (ms)** | **MNI coordinates** |  |  | **T (9)** | **Effect size (s.e.m.)** |
| --- | --- | --- | --- | --- | --- | --- | --- |
| Cat>Val | mSTG-R* | 157 | 66 | -30 | 10 | 5.30 | 2.24(0.05) |
| Cat>Val | mSTG-R | 198 | 66 | -22 | 4 | 5.26 | 1.53(0.04) |
| Cat>Val | aSTS-R | 357 | 56 | 0 | -13 | 3.81 | 0.62(0.02) |
| Cat>Val | pSTG-R | 357 | 56 | -52 | 22 | 3.63 | 0.47(0.01) |
| Cat>Aro | mSTG-R* | 157 | 66 | -37 | 4 | 5.11 | 2.56(0.07) |
| Aro>Cat | mSTG-R | 717 | 66 | -30 | 2 | 4.00 | 2.73(0.08) |
| Aro>Cat | IFGop-R* | 717 | 51 | 0 | 20 | 4.08 | 2.22(0.03) |
| Aro>Val | mSTG-R* | 637 | 61 | -10 | -3 | 4.64 | 5.61(0.20) |

**Table S3:**
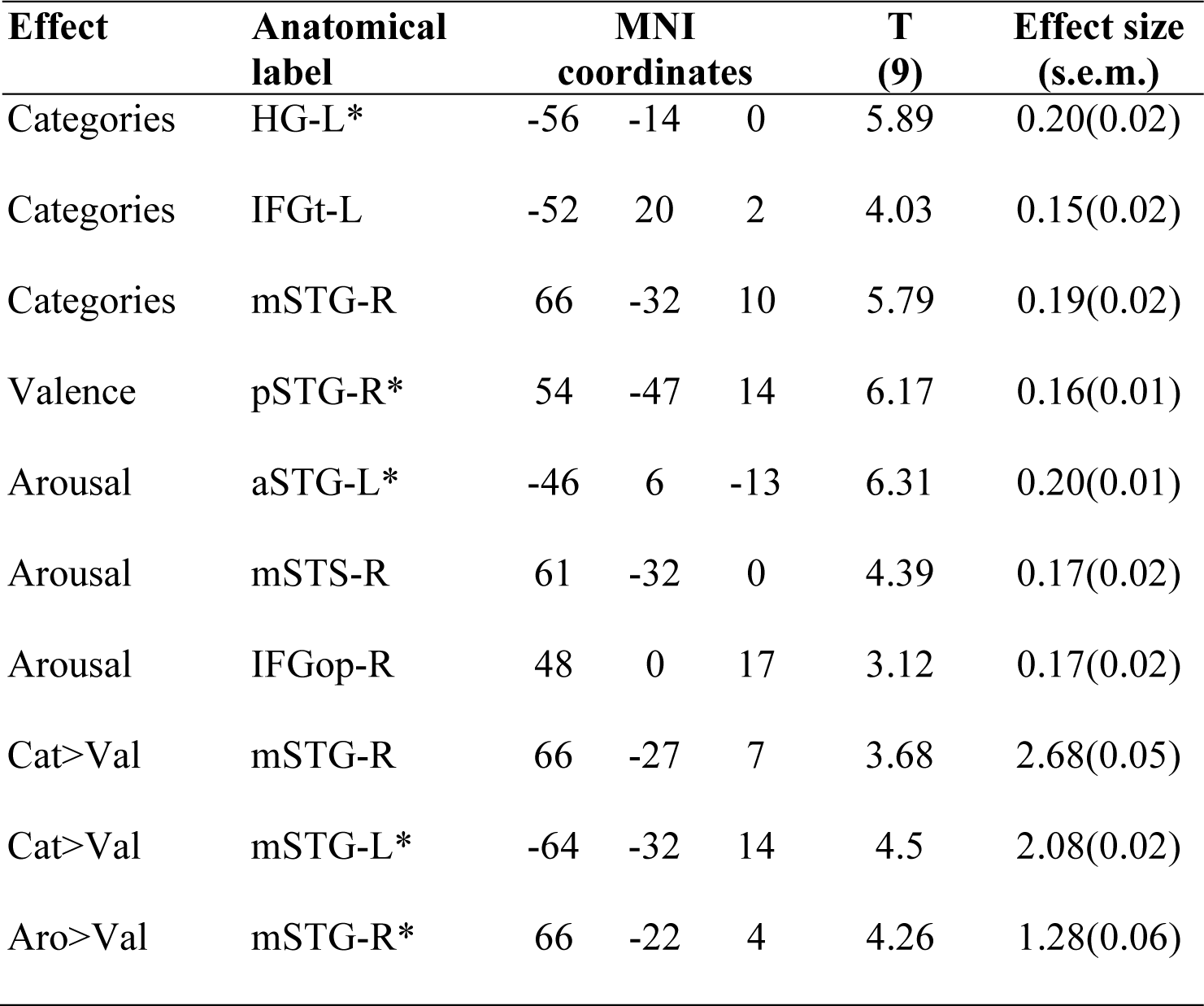
**Global and local peaks of emotion encoding and explained variance contrast analyses in fMRI data.** The table lists global and local peaks in the GLM T-maps. HG-L = left Heschl’s gyrus; IFGt/op = inferior frontal gyrus, pars triangularis/opercularis; m/a STG/STS = mid/anterior superior temporal gyrus/sulcus. All effects significant at p < 0.05 FWE corrected within the analysis mask. See Table S1 for further details.

| **Effect** | **Anatomical label** | **MNI coordinates** |  |  | **T (9)** | **Effect size (s.e.m.)** |
| --- | --- | --- | --- | --- | --- | --- |
| Categories | HG-L* | -56 | -14 | 0 | 5.89 | 0.20(0.02) |
| Categories | IFGt-L | -52 | 20 | 2 | 4.03 | 0.15(0.02) |
| Categories | mSTG-R | 66 | -32 | 10 | 5.79 | 0.19(0.02) |
| Valence | pSTG-R* | 54 | -47 | 14 | 6.17 | 0.16(0.01) |
| Arousal | aSTG-L* | -46 | 6 | -13 | 6.31 | 0.20(0.01) |
| Arousal | mSTS-R | 61 | -32 | 0 | 4.39 | 0.17(0.02) |
| Arousal | IFGop-R | 48 | 0 | 17 | 3.12 | 0.17(0.02) |
| Cat>Val | mSTG-R | 66 | -27 | 7 | 3.68 | 2.68(0.05) |
| Cat>Val | mSTG-L* | -64 | -32 | 14 | 4.5 | 2.08(0.02) |
| Aro>Val | mSTG-R* | 66 | -22 | 4 | 4.26 | 1.28(0.06) |

**Table S4:**
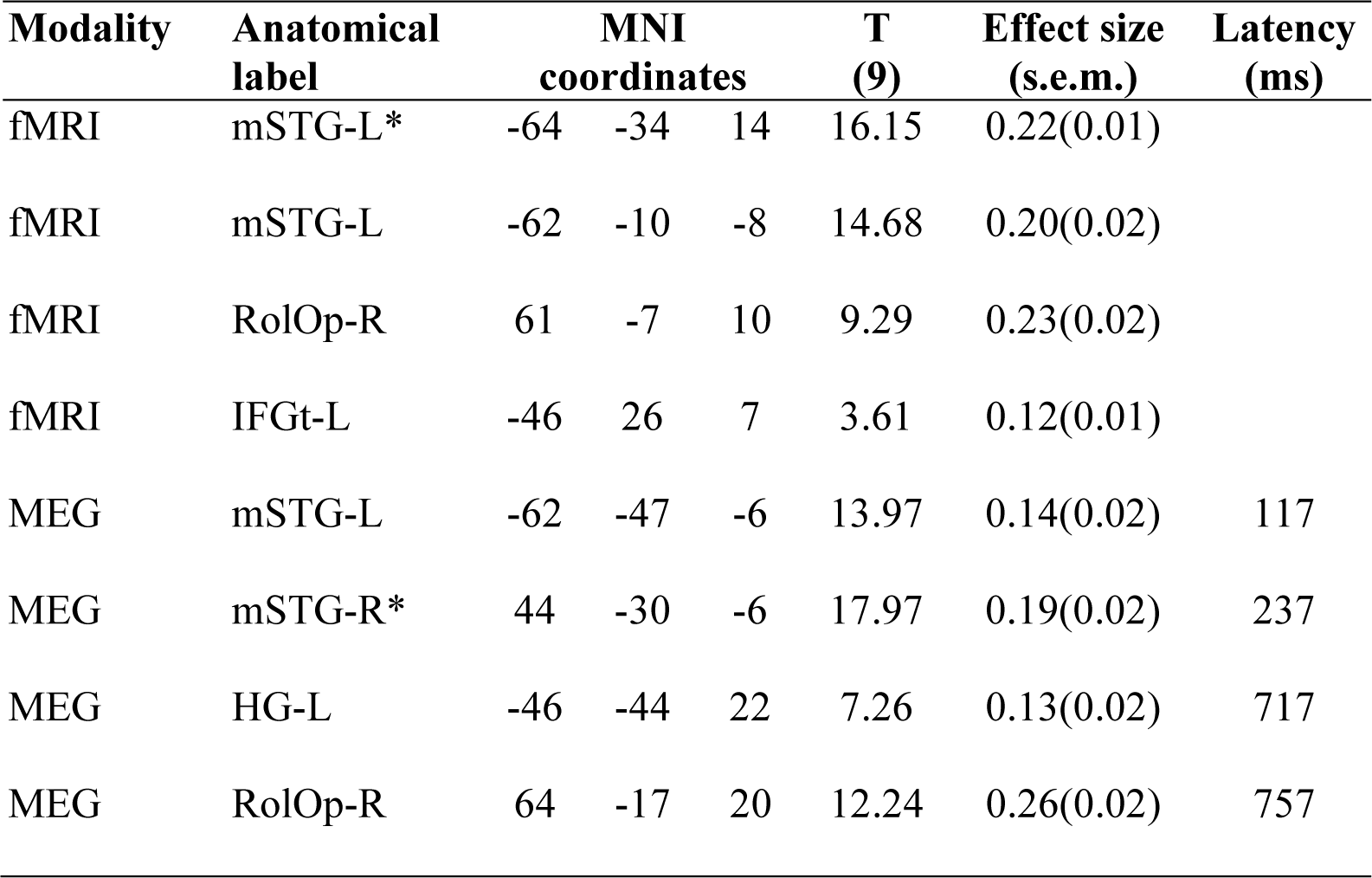
**Global and local peaks of encoding of perceived voice dissimilarity in fMRI and MEG cerebral response patterns.** The table lists global and local peaks in the GLM T-maps for testing the correlation between fMRI and MEG cerebral response patterns and perceptual voice dissimilarity as estimated from behavioral responses. mSTG-L = right mid superior temporal gyrus; RolOp = Rolandic operculum; HG-L = left Heschl’s gyrus. All effects significant at p < 0.05 FWE corrected within the analysis mask. See Table S1 for further details.

